# Bottom trawling impacts on faunal and microbial communities increase seafloor carbon loss

**DOI:** 10.64898/2026.01.29.702536

**Authors:** Guido Bonthond, Jan Beermann, Alexa Wrede, Vera Sidorenko, Peter J. Schupp, Lars Gutow

## Abstract

The seafloor is the largest organic carbon sink on Earth. Its benthic fauna and microbiota jointly regulate long-term carbon storage by assimilating and degrading sedimentary organic matter (SOM). Bottom trawling threatens the long-term storage of organic matter by physical disturbance of the seafloor and remobilization of SOM. Trawling also impacts faunal and microbial communities and their diversity, but the implications of these ecological consequences for long-term carbon storage dynamics are poorly understood.

Here, we combined datasets of paired faunal and microbial communities from the North Sea and used structural equation modeling to disentangle direct and biotically mediated effects of bottom trawling on SOM. One third of trawling-induced SOM loss is indirect and mediated by shifts in benthic faunal and microbial communities. Microbial community structure directly affected SOM and represented a key link in this pathway. Microbial effects on SOM were strongly tied to changes in metabolic traits. SOM loss coincides with increased microbial aerobic respiration potential, whereas SOM increases correlated with microbial assimilation processes, including dark carbon fixation. Benthic fauna impact SOM indirectly by shaping benthic microbial communities. Faunal diversity and species’ traits related to bioturbation promoted shifts in benthic microbiota that favor OM storage, corroborating their indirect but essential role in seafloor carbon dynamics.

This study reveals that bottom trawling alters seafloor carbon storage not only through physical disturbance but also by disrupting faunal-microbial interactions that regulate sediment carbon dynamics. Taxonomic and functional shifts in benthic microbiota are a key, but underestimated, component of the processes regulating the seafloor carbon sink and warrant better integration of benthic biodiversity and ecological processes into assessments of seafloor carbon storage under anthropogenic pressures.

## INTRODUCTION

Marine sediments cover nearly 70% of Earth’s surface and critically contribute to the long-term storage of organic matter (OM), of which organic carbon (OC) is the major constituent (Snelgrove *et al*. 2014; Atwood *et al*. 2020). Sedimentary organic matter (SOM) originates from marine primary production (phytoplankton and macrophytes) and terrestrial runoff that is deposited as detritus on the seafloor. This detritus is degraded by benthic invertebrates and microbes, but a fraction escapes heterotrophic consumption and may be buried and stored in the seafloor for thousands to millions of years (Estes *et al*. 2019; Atwood *et al*. 2020). As the ultimate decomposers of OM, benthic invertebrates and microbial heterotrophs jointly regulate the storage of OM in the seafloor and thereby play a key role in the carbon cycle.

Carbon sequestration is especially pronounced in productive coastal shelf seas, globally accounting for more than 85% of all OC that is annually buried in marine sediments (Atwood *et al*. 2020; Epstein *et al*. 2022). At the same time, shelf seas are most exposed to multiple human impacts, such as pollution, sand extraction, mining, infrastructure construction, and bottom trawling by demersal fisheries (Amoroso *et al*. 2018; Coates *et al*. 2014; Harris 2020; Hiddink *et al*. 2017; Porz *et al*. 2026), which may affect the capacity of sediments for OC sequestration (Epstein *et al*. 2022). In particular, bottom trawling impacts benthic ecosystems by altering seabed morphology (Puig *et al*. 2012) and resuspending large quantities of sediment (Breimann *et al*. 2022), thereby reducing OC storage in exchange for carbon dioxide emissions (Bradshaw *et al*. 2024; Tiano *et al*. 2024; Zhang *et al*. 2024). Furthermore, trawling causes high mortality in benthic macrofauna (Tillin *et al*. 2006; Hiddink *et al*. 2017; Sciberras *et al*. 2018) and alters benthic microbial communities and metabolism (Bruce *et al*. 2022; Bonthond *et al*. 2023), which may even be detectable in microbial communities of seafloor-inhabiting holobionts, such as flatfish (Gwinner *et al*. 2026).

The impact of bottom trawling on seafloor carbon storage is a matter of growing concern. Trawling promotes the degradation of the more reactive fraction of SOC, but its effect on total SOC is context-dependent (Porz *et al*. 2024; Tiano *et al*. 2024; Zhang *et al*. 2024). While mechanical impacts (i.e., heavy gear disturbing the sediment) directly remobilize OC, the depletion of benthic fauna can also have consequences for SOC (Epstein *et al*. 2022), as these organisms degrade and assimilate detritus and actively rework the sediment matrix through bioturbation. Bioturbation broadly refers to all faunal transport processes within the sediment, including particle reworking and burrow ventilation (Kristensen *et al*. 2012). Bioturbating macrofauna also influence benthic microbial communities by transporting oxygen and OC to deeper sediment layers, and by homogenizing benthic microbiota organized along redox gradients (Deng et al. 2020; Middelburg 2018). Accordingly, trawling-induced shifts in microbial diversity and metabolism (Bruce *et al*. 2022; Bonthond *et al*. 2023) may be partially mediated by changes in macrofaunal communities.

Benthic microbiota in turn degrade and assimilate detritus and therewith impact the storage and cycling of OC (Jiao *et al*. 2010; LaRowe *et al*. 2020). However, the roles of microbes in SOM cycling, and how their interactions with benthic macrofauna influence SOM, are not well understood (Jiao *et al*. 2010; Middelburg 2018; LaRowe *et al*. 2020). In terrestrial environments, microbiota are known to critically impact the accumulation and degradation of OC in soils. Due to their capacity to decompose recalcitrant OC, microbial respiration can enhance OC remineralization and reduce soil OC (De Graaff *et al*. 2015). At the same time, microbial biomass and byproducts promote OC stabilization and sequestration (Derrien *et al*. 2023; Tao *et al*. 2023). If microbial processes are of similar importance for OC storage in the seafloor, the interplay among macrofauna and microbiota could importantly mediate anthropogenic impacts, such as bottom trawling, on seafloor carbon storage.

Here, we used an extensive paired dataset of benthic macrofaunal and microbial communities to disentangle direct and indirect effects of bottom trawling on SOM. One hundred and fifty sites were sampled across a regional scale in the southeastern North Sea, a region subject to variable fishing intensities and characterized by a broad range of SOM contents. We used structural equation models (SEMs) to analyze how SOM is jointly shaped by abiotic factors (sediment characteristics, temperature, shear stress, trawling) and by macrofaunal and microbial communities. We hypothesized that benthic macrofauna and microbiota mediate the effects of bottom trawling on SOM. To test this hypothesis, we developed a conceptual framework including different potential abiotic and biotic pathways through which trawling may influence SOM. Further, we used multivariate and penalized generalized linear models to identify key faunal and microbial traits and taxa, responding to trawling, and with impact on SOM.

## RESULTS

### Sampling and data description

Macrofaunal and microbial communities were characterized for 150 sites in the southeastern North Sea (Figure 1). Sediment organic matter contents varied from 0.063 to 5.02% weight and trawling intensities (in swept area ratio per year, SAR y^-1^) from close to zero (0.016 SAR y^-1^) to heavy trawling rates of up to 1.262 SAR y^-1^.

**Figure 1.**
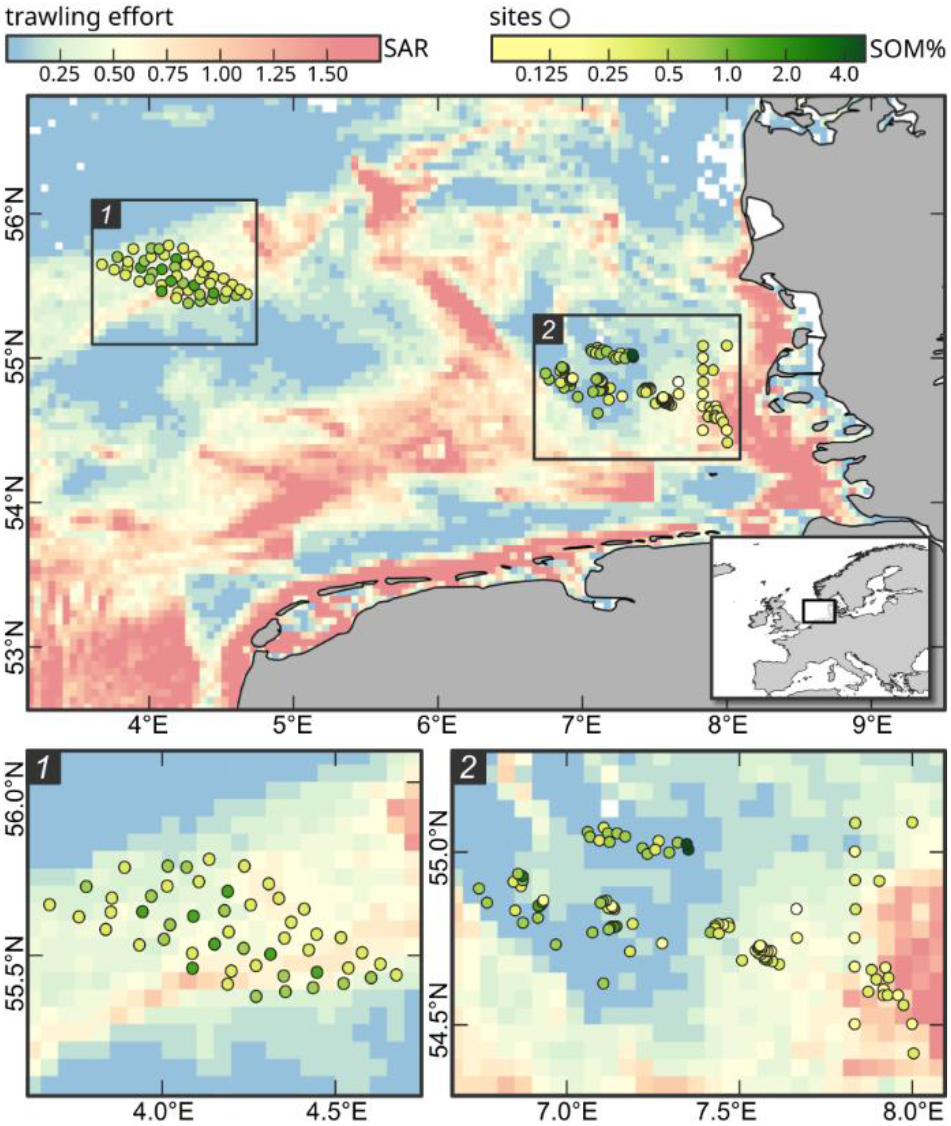
Geographic overview of all study sites in the southeastern North Sea where macrofaunal and microbial samples were obtained in parallel. The background heatmap displays the bottom trawling intensity calculated as swept area ratio (SAR) per year.

### Benthic microbiota

The microbial dataset counted 69,556 OTUs with 1,293 different genus classifications. Non-metric multidimensional scaling (NMDS) revealed various trends associated with microbial community structure (Figure 2A). Apart from shear stress and microbial diversity, all variables were significantly ordinated on the NMDS space (p < 0.05, Table S1). SOM increased along the second NMDS axis, while trawling intensity decreased (Figure 2B-C). Microbial community structure also varied with faunal community properties. The biomass, the community bioturbation potential (BPc, Queirós *et al*. 2013), and the community bioirrigation potential (IPc, Wrede *et al*. 2018) were ordinated in similar directions, increasing along NMDS1 and NMDS2. Faunal diversity decreased along both axes.

**Figure 2.**
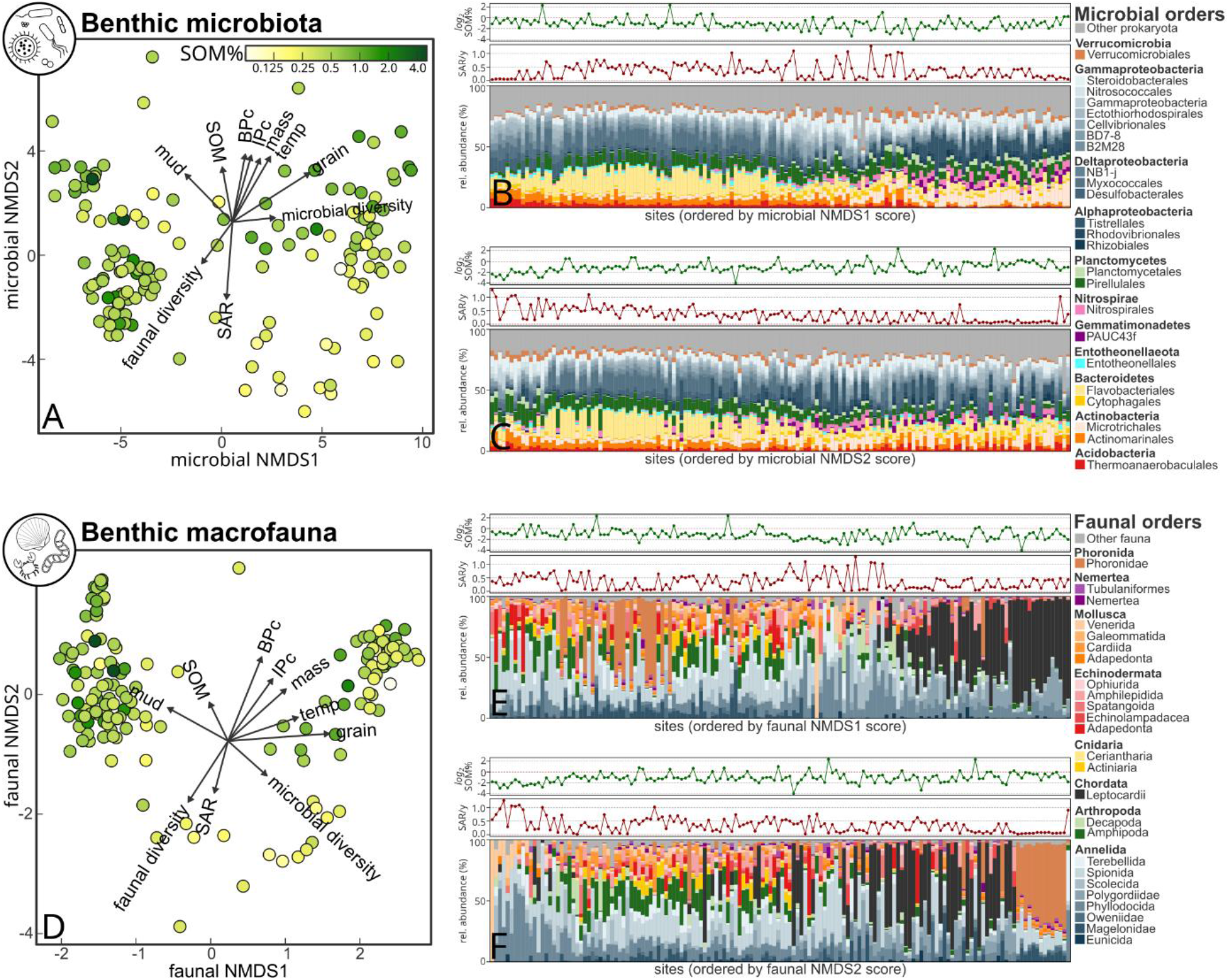
Community composition of benthic microbiota **(A-C)** and macrofauna **(D-F)** across 150 stations. **(A, D)** NMDS ordination plots based on **(A)** benthic microbial community composition using Aitchison distances (stress = 0.097), and **(D)** benthic faunal community composition using Bray-Curtis distances (stress = 0.128). Predictor and response variables fitted on the ordination are indicated by arrows (showing only variables with p < 0.05). **(B-C, E-F)** Stacked barplots showing the 25 relative most abundant microbial **(B-C)** and faunal **(E-F)** orders of all 150 sites, ordered by their scores on the respective NMDS axes. Border plots on top of each barplot show the log transformed SOM% (green), and trawling intensity in SAR y^-1^ (red).

Pseudomonadota was the most abundant phylum, followed by Bacteroidetes and Planctomycetes. Compositional changes along NMDS1 included a shift within the Pseudomonadota from Delta-to Gammaproteobacteria. Bacteroidetes decreased along NMDS1 while Nitrospirae increased. Actinobacteria transitioned from Actinomarinales to Microtrichales (Figure 2B). Compositional changes along NMDS2 were characterized by a strong decrease in Planctomycetes and an increase in Nitrospirae, but were otherwise less pronounced at higher taxonomic ranks (Figure 2C).

### Benthic macrofauna

In total, 301 faunal taxa were identified, of which 218 taxa were classified to species and 39 to genus level. Besides bottom shear stress, all variables were significantly ordinated onto the NMDS space (p < 0.05, Figure 2D). Compositional changes along NMDS1 coincided with a decrease in mud content and an increase in temperature and median grain size. Along NMDS2, SOM increased and trawling intensity decreased (Figure 2F). Faunal biomass, BPc, and IPc also increased along NMDS2. Faunal diversity decreased along both NMDS axes, and microbial diversity increased along NMDS1 and decreased along NMDS2.

Annelida were the largest and most diverse phylum (36.7% relative abundance, 115 taxa, Figure 2E-F). Phoronida and Chordata were also among the most abundant phyla (24.0% and 15.2% relative abundance), but counted only two taxa each. NMDS1 showed a strong shift within the Annelida from Spionida to Phyllodocida and Polygordiidae, and to communities dominated by *Branchiostoma lanceolatum* (Chordata, Figure 2E). Along NMDS2, faunal community composition gradually shifted from communities dominated by Annelida, along high abundances of both Annelida and Arthropoda, toward communities dominated by *Phoronis muelleri* (Phoronidae, Figure 2F).

### Structural equation modeling of abiotic and biotic effects on microbial composition and SOM

The initial SEM did not support faunal biomass and the IPc as informative predictors for microbial community composition or SOM. Accordingly, they were excluded from the final model, resulting in a better fit (Fisher’s C = 51.893, df = 42, p = 0.141, AICc = 4433.95, Table S2-4). Median grain size, mud content, and trawling intensity had direct effects on faunal community composition. Faunal diversity was directly dependent on faunal community composition, increased with temperature, and decreased with trawling intensity (Figure 3A). Direct abiotic effects on microbial community structure were in descending order of strength: median grain size, mud content, and trawling intensity. Microbial community structure was also dependent on faunal composition, the BPc, and faunal diversity. Consequently, fauna mediated the effects of temperature, mud content, median grain size, and trawling intensity on microbial composition. These indirect effects were substantial and equaled about half of the direct effect in case of mud content and median grain size. For trawling intensity, the indirect effect on benthic microbial composition was even larger than the direct effect (Figure 3B). Microbial diversity varied with microbial and faunal composition, but not significantly with faunal diversity. Shear stress was the only predictor for which no effects were statistically supported.

**Figure 3.**
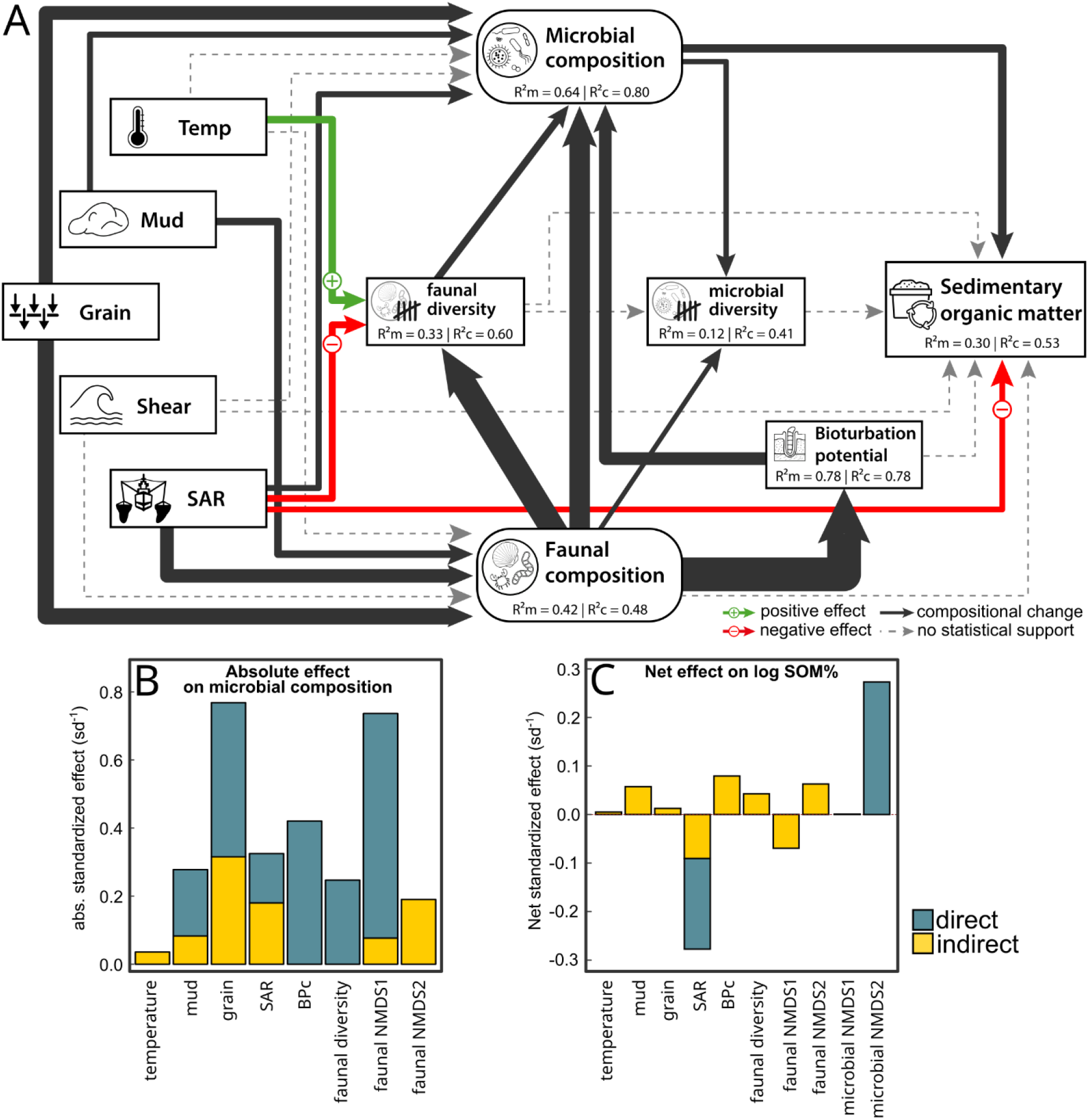
**(A)** Structural equation model of SOM in response to abiotic and biotic variables. Red and green arrows indicate significant (p < 0.05) negative and positive effects. Black arrows indicate significant effects associated with changes in composition and dashed gray arrows show hypothesized relationships which were statistically not supported. Arrows widths are proportional to their standardized effect sizes. Observed variables are shown in squared boxes and latent variables in rounded boxes. The marginal and conditional R^2^ values of the LMMs are shown in boxes of the respective response variables, indicating the model variation explained by fixed effects (R^2^m), and by both fixed and random effects (R^2^c). **(B)** Absolute direct (blue) and indirect (yellow) standardized effects on microbial community composition. Displayed values are the sum of the significant (p < 0.05) standardized effects of the same variable on microbial NMDS1 and NMDS2. **(C)** Net direct (blue) and indirect (yellow) standardized effects of log transformed SOM%.

Bottom trawling effort directly reduced SOM. Additionally, the microbial latent variable NMDS2 was significant, indicating a direct effect of microbial community structure on SOM (Figure 3A). The effect of microbial diversity on SOM was not significant, and neither were faunal composition, BPc and faunal diversity. Several pathways were supported through which faunal composition, BPc and faunal diversity influenced SOM indirectly. The indirect effects of faunal diversity and BPc on SOM where both positive, whereas the indirect effect of trawling was negative (Figure 3C).

### Bioturbation related traits and taxon level effects

Trawling impacted in total 74 predicted pathways (Figure 4A, Figure S3A), which were mostly related to biosynthesis (13 decreasing, 64 increasing). In addition, especially pathways related to energy metabolism (belonging to the categories ‘generation of precursor metabolites’ and ‘degradation/utilization/assimilation)’ increased in response to trawling (6 and 5, respectively), including the TCA cycle I (*TCA*) and VI (*PWY-5913*), and aerobic respiration I (*PWY-3781*). Various OTUs varied with trawling effort (Figure 4B). The relative abundance of OTUs within the Actinobacteria and Planctomycetes, particularly increased with trawling effort, whereas Alphaproteobacteria and Deltaprotebacteria tended to decrease. Fifteen faunal traits declined with trawling effort and none increased (Figure 4C), suggesting an overall decrease in faunal functional diversity. This was also reflected at the species level, where the abundance of 27 species decreased and only 12 species increased in response to trawling (Figure 4D). In particular, species belonging to Annelida, Mollusca and Nemertea decreased with trawling intensity, whereas in other phyla increasing and decreasing taxa were more balanced.

**Figure 4.**
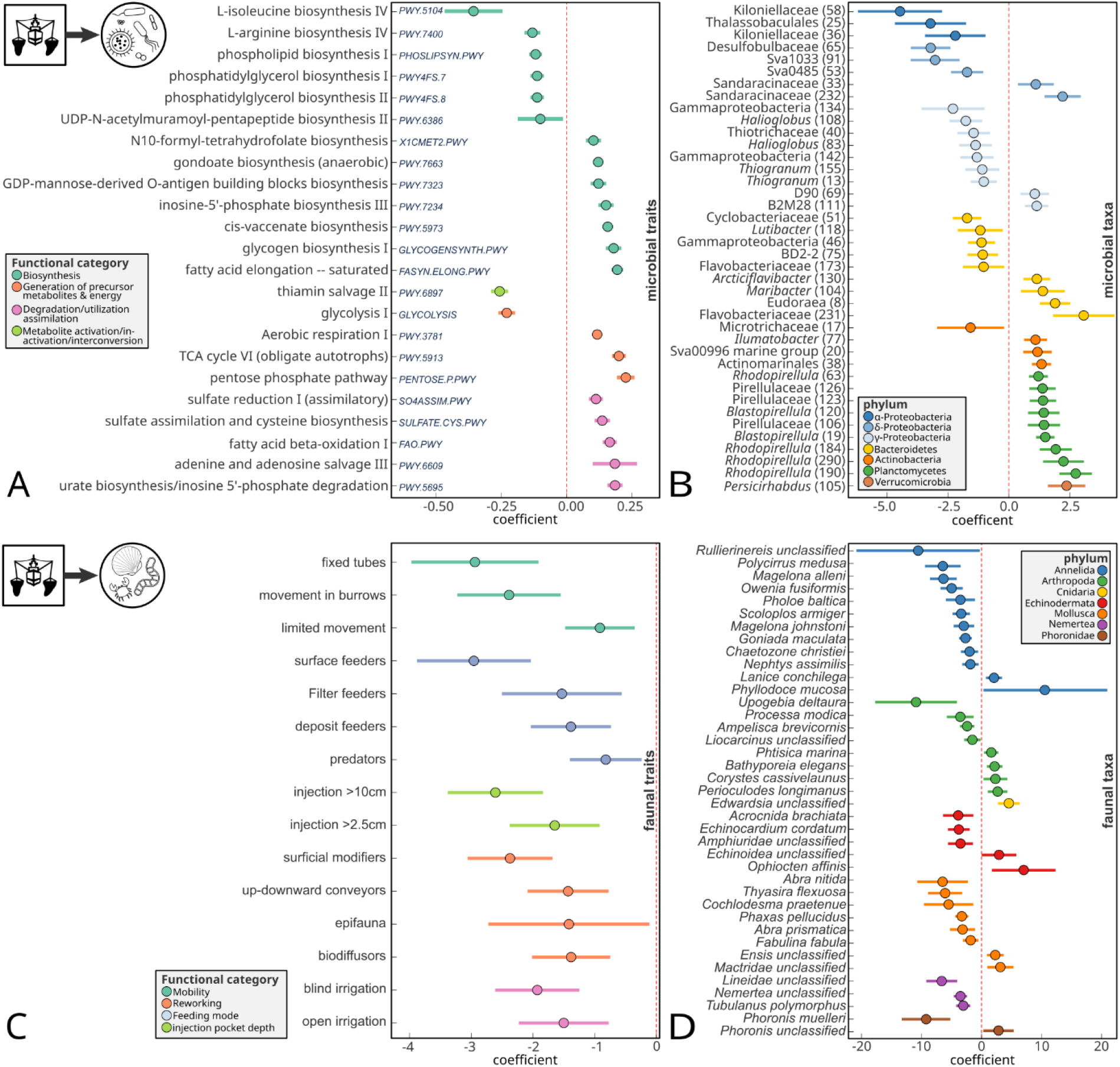
Trawling impacts on microbial **(A, B)** and faunal **(C, D)** traits and species identified through mGLMs. (A) Microbial traits (predicted pathways) with significant responses to trawling effort and effect sizes > 0.1. **(B)** Microbial taxa (OTU numbers in brackets) with significant responses to trawling effort and effect sizes > 0.1. **(C)** Faunal traits with significant responses to trawling effort. **(D)** Faunal taxa with significant responses to trawling effort. Colors correspond to MetaCyc functional categories **(A)**, faunal trats **(C)** phylum-level taxonomy **(B, D)**. Se supplementary 1 for the extended figure including micorbial traits and taxa with smaller effect sizes.

In total, eight trait categories were identified with pGLMs to explain changes in microbial community structure (Figure 4A). Most of these traits were related to mobility (limited movement, free movement through sediment matrix, free movement through burrows) and reworking (bio-diffusors, up- and downward conveyors, surficial modifiers), but also included the feeding type category ‘subsurface filter feeders’, and the injection pocket depth category ‘below 10 cm’. The model retained all environmental covariates (i.e., mud content, median grain size, trawling intensity), the faunal latent variables, and faunal diversity. When analyzing taxa instead of traits, 30 faunal taxa were identified to shape microbial community composition (Figure 4B). More than half of these taxa belonged to the Annelida, with the two species *Eteone longa* and *Aonides paucibranchiata* having the highest impact. But also *B. lanceolatum* (Chordata) and *Echinocardium pennatifidum* (Echinodermata) were among the species with highest effect scores.

Twenty-seven predicted microbial pathways were resolved as informative predictors of SOM (Figure 4C). Eighteen pathways were negatively associated with SOM, of which the strongest effect was observed for aerobic respiration (*PWY-3781*). Out of the nine pathways associated with an increase in SOM, the two strongest regularized coefficients were found for gluconeogenesis (*GLUCONEO-PWY*) and the Calvin cycle (*CALVIN-PWY*). Twenty-five OTUs were associated with SOM decrease, and four OTUs with SOM increase (Figure 4D). Planctomycetes were prominent among the OTUs related to SOM decrease (nine in total), but the strongest negative association was found for an Actinobacterial OTU, classified to Sva0996 marine group. From the covariates, trawling intensity, microbial NMDS2, and predicted functional microbial diversity were retained in both pGLMs evaluating trait and species effects on SOM.

## DISCUSSION

Our study confirms that bottom trawling reduces OC storage in marine sediments. While the trawling impact is partially direct, likely resulting from physical seabed disturbance, approximately one third of the overall bottom trawling impact on SOM is indirect, mediated by changes in benthic macrofaunal and microbial communities. Our analysis provides insight into these indirect trawling effects, and reveals several pathways through which coexisting fauna and microbiota shape the sequestration of carbon. The direct link between benthic microbial community structure and SOM is central for these pathways. Consequently, benthic microbiota serve as a key mediator of various biotic and abiotic effects on OC storage. While macrofaunal communities could not be directly linked with SOM, they strongly impact microbial composition, marking an indirect, but pivotal, effect of macrofauna on SOM. The net effects of faunal diversity and bioturbation on SOM are positive. These results indicate that the indirect impacts of bottom trawling on SOM, cascading through macrofaunal and microbial communities, have net negative consequences for the seafloor carbon stocks. Together, our findings emphasize the central role of benthic fauna and microbiota, and the interaction among them, in OC sequestration and underscore the need to consider both physical and ecological processes for the assessment of long-term consequences of bottom trawling on seafloor carbon dynamics.

### Trawling reduces SOM by restructuring benthic fauna and microbiota

The impact of bottom trawling on OM storage in sediments entails a combination of mechanisms (Epstein *et al*. 2022; Zhang *et al*. 2024). The mechanical resuspension of OM is the most direct effect, as it locally mobilizes OM from the sediment, which can then become subject to export by currents, and is partially remineralized by heterotrophs before being deposited, and buried again. In addition, physical remixing of the upper sediment layers enhances benthic remineralization of OM in the sediment through aerobic microbial respiration. Previous studies have demonstrated that the extended exposure to oxygen, caused by resuspension and remixing, decreases SOM, providing estimates of trawling induced loss of OC stored in the seafloor (Sala *et al*. 2021; Hiddink *et al*. 2023; Porz *et al*. 2024; Zhang *et al*. 2024). Our results confirm the negative direct impact of trawling on SOM. Additionally, our study reveals that trawling-induced shifts in benthic macrofaunal and microbial communities also have important consequences for carbon storage, supporting our hypothesis that macrofauna and microbiota mediate the effects of bottom trawling on SOM. Trawling alters the communities of organisms involved in the processing and degradation of SOM. The contribution of benthic biota to SOM dynamics has been acknowledged in previous studies (Epstein *et al*. 2022; Zhang *et al*. 2024). However, the biotic pathways involved have been poorly characterized and are not explicitly considered in predictive models of sedimentary carbon storage. Instead, these models typically estimate SOM remineralization as a function of extended oxygen exposure associated with trawling-induced sediment disturbance, treating the benthic biota rather as a ‘black-box’ without resolving the contribution of the fauna and microbiota mechanistically (Sala *et al*. 2021; Zhang *et al*. 2024). Our analysis, however, demonstrates that indirect, biota-mediated effects are crucial and complex, accounting for roughly one third of the total trawling impact on SOM (Figure 3C).

Benthic microbiota were identified as a key mediator of trawling impacts on SOM. Our model showed that compositional shifts in benthic microbiota are directly linked to changes in SOM, highlighting the prominent role of microbial processes in shaping SOM. Bottom trawling induces changes in the composition and energy metabolism of benthic microbial communities promoting specifically aerobic metabolic pathways (Bonthond *et al*. 2023). The responses to trawling varied between microbial groups with some groups apparently benefiting from trawling while other groups clearly declined in response to trawling. The study confirmed that the predicted aerobic respiration pathway increased with trawling effort, indicating that microbial aerobic heterotrophic metabolism is more prevalent in sediments exposed to intense trawling. These taxonomic and functional shifts in the microbial communities explain a net loss in SOM, highlighting microbial aerobic respiration as the critical process by which SOM is removed from the food web (Middelburg 2018).

The impacts of trawling on benthic fauna also have negative consequences for seafloor carbon storage. In line with previous research (see Sciberras *et al*. 2018 and references therein), various traits related to bioturbation also benthic faunal diversity reduced with trawling intensity. This trawling-induced decline in structural and functional diversity of the benthic macrofauna coincided with, but did not directly cause, a decrease in SOM. Instead, trawling induces a cascade of responses involving both benthic macrofauna and microbiota. Losses in response to trawling were particularly pronounced in annelids and mollusks and especially these groups impacted microbiota. As macrofaunal diversity and bioturbation regulate benthic microbiota, there is an indirect negative net effect of trawling on SOM, mediated by the benthic fauna. Accordingly, marine protection efforts to enhance benthic faunal diversity have the potential to also promote seafloor carbon storage.

### Carbon storage is directly affected by benthic microbiota

Aerobic respiration is the terminal process that catalyzes the conversion of OC to carbon dioxide (Middelburg 2018; Jørgensen *et al*. 2022). In the pelagic environment, OC is primarily recycled through the microbial loop, where OC assimilated by microbes is partially transferred to higher trophic levels through predation and grazing (Azam *et al*. 1983). The transfer of microbially processed OC to higher trophic levels in sediments is, however, limited and most OC consumed by microbes is eventually lost through microbial respiration, a concept described as the ‘inverted microbial loop’ (Middelburg 2018). In accordance with this, microbial aerobic respiration emerged in our analysis as the most important pathway associated with SOM loss. In Bonthond *et al*. (2023) and in this study, we found that the predicted aerobic respiration pathway increased with trawling effort, indicating that microbial aerobic heterotrophic metabolism is more prevalent in sediments exposed to more trawling. Both macrofauna and microbes carry out aerobic respiration. However, they partially utilize different OM substrates from different sources. This likely determines whether SOM is being decomposed and cycled back into the food web through assimilation, or whether it escapes degradation and becomes sequestered. Living tightly attached to sand grains, microbes utilize sediment-bound particulate OM (Arnosti 2011). This OM is characterized by its low reactivity with the labile OM fraction typically accounting for <10% (Pusceddu *et al*. 2009; Arndt *et al*. 2013). Among prokaryotes, sophisticated metabolic pathways have evolved, enabling the degradation of even the most recalcitrant molecules (Jiao *et al*. 2010; LaRowe *et al*. 2020), which are stable and most likely to be sequestered. Essentially, under adequate conditions, there are no biomolecules known that cannot be degraded by microorganisms (LaRowe *et al*. 2020; Dittmar *et al*. 2021). One of the phylum-level microbial shifts associated with decreasing SOM was a relative increase in Planctomycetes, which include microbial groups specialized in decomposing high molecular weight and diverse recalcitrant OM substrates (Probandt *et al*. 2017; Wiegand *et al*. 2018). Additionally, Planctomycetes were prevalent among the OTUs that increased with trawling effort, indicating that bottom trawling not only promotes OM remineralization by enhancing oxygen exposure, but also by favoring microbial metabolic degradation of less reactive OM, which is more probable to be sequestered. Thus, trawling not only enhances the exposure of SOM to oxygen, but also to microbial decomposers capable of remineralizing low-reactive OM.

Our analysis identified the Calvin cycle as a primary predicted microbial function associated with SOM increase. As light availability at the sampled sites is negligible to absent, this likely reflects dark carbon fixation by chemoautotrophic microbes, a common process in marine sediments, largely driven by Gammaproteobacteria (Hügler & Sievert 2011; Middelburg 2011; Dyksma *et al*. 2016). The strong increase of SOM with the gluconeogenesis pathway is further in line with this. Gluconeogenesis is an assimilatory pathway, which synthesizes glucose from smaller precursors, such as fatty acids and amino acids, and indirectly from carbon dioxide (Sharkey & Weise 2012). Dark carbon fixation in sediments is recognized as an important contribution to the input of new OC into the oceans and its burial in sediments (Middelburg 2011), but has not been considered in the context of bottom trawling. Accordingly, trawling may not only enhance the remineralization of OC but also reduce the benthic OC production by chemoautotrophs.

The weak positive effect of the predicted functional microbial diversity on SOM suggests that degradation and accumulation of SOM involves ‘diversity effects’. Diversity effects define contributions of a community to processes that cannot be explained by the sum of the contribution of its single members, and are known to play an important role in OC storage in terrestrial ecosystems (Gessner *et al*. 2010). The decomposition of organic substrates involves complex trophic interactions, as well as competition and cooperation among primary degraders of recalcitrant molecules (e.g., cellulose, lignin, laminarin) and secondary consumers of more labile OM (e.g., acetate, lactate, amino acids; Gralka *et al*. 2020). Functional diversity and redundancy influence these interactions and increase carbon use efficiency, potentially promoting OC respiration and decomposition with negative impact on OM storage (De Graaff *et al*. 2015). At the same time, the increased generation of microbial biomass and more stable by-products can promote OC accumulation (Tao *et al*. 2023). Our results indicate that microbial functional diversity influences OC cycling in marine sediments in a similar manner to terrestrial soils. This is of particular importance in the context of bottom trawling impacts on seafloor carbon dynamics, which can decrease microbial taxonomic and functional diversity (Bonthond *et al*. 2023).

### Faunal communities influence SOM by regulating benthic microbiota

Faunal composition, bioturbation, and diversity had no direct effect on SOM but significantly influenced microbial community structure, highlighting that benthic microbial diversity and community composition are shaped not only by abiotic factors (Bonthond *et al*. 2023), but also by the composition of macrofaunal communities. Consequently, macrofauna mediate the effects of different environmental processes on benthic microbiota, constituting a key link between bottom trawling and carbon sequestration.

A direct effect of bioturbation on SOM was not supported, suggesting only minor effects of sediment reworking on SOM. Alternatively, antagonistic processes inherent to bioturbation, such as up- and downward OC transport, may minimize the net effect of bioturbation. Our results indicate that bioturbation essentially modulates benthic microbiota. This is in agreement with previous work, that demonstrated that faunal bioturbation activity shapes benthic microbiota by vertically mixing OC substrates and homogenizing microbial communities (Deng *et al*. 2020). The BPc, which is a commonly used index to estimate the community bioturbation potential (Queirós *et al*. 2013), explained shifts in benthic microbiota. Specifically, faunal traits related to mobility and sediment reworking activity, which are used to calculate the BPc, were identified as important predictors of SOM (Figure 5A) and also declined with trawling intensity (Figure 4C). Bioturbators have been shown to promote aerobic microbial OC degradation by enhancing oxygen supply to the sediment and its microbiota (Baranov *et al*. 2016). At the same time, macrofauna compete with the microbiota for oxygen (Middelburg 2018; Jørgensen *et al*. 2022), which may locally limit microbial aerobic respiration. The strong impact of the faunal trait ‘limited movement’ on benthic microbiota may indicate a suppression of microbial aerobic respiration by stationary animals that consume oxygen within the sediment but enhance its resupply only minimally. Additionally, benthic fauna may contribute to the stabilization of SOM as their feces and remains contain more recalcitrant OC compared to freshly deposited detritus and are therefore degraded less efficiently by microbes (Arndt *et al*. 2013; LaRowe *et al*. 2020).

**Figure 5.**
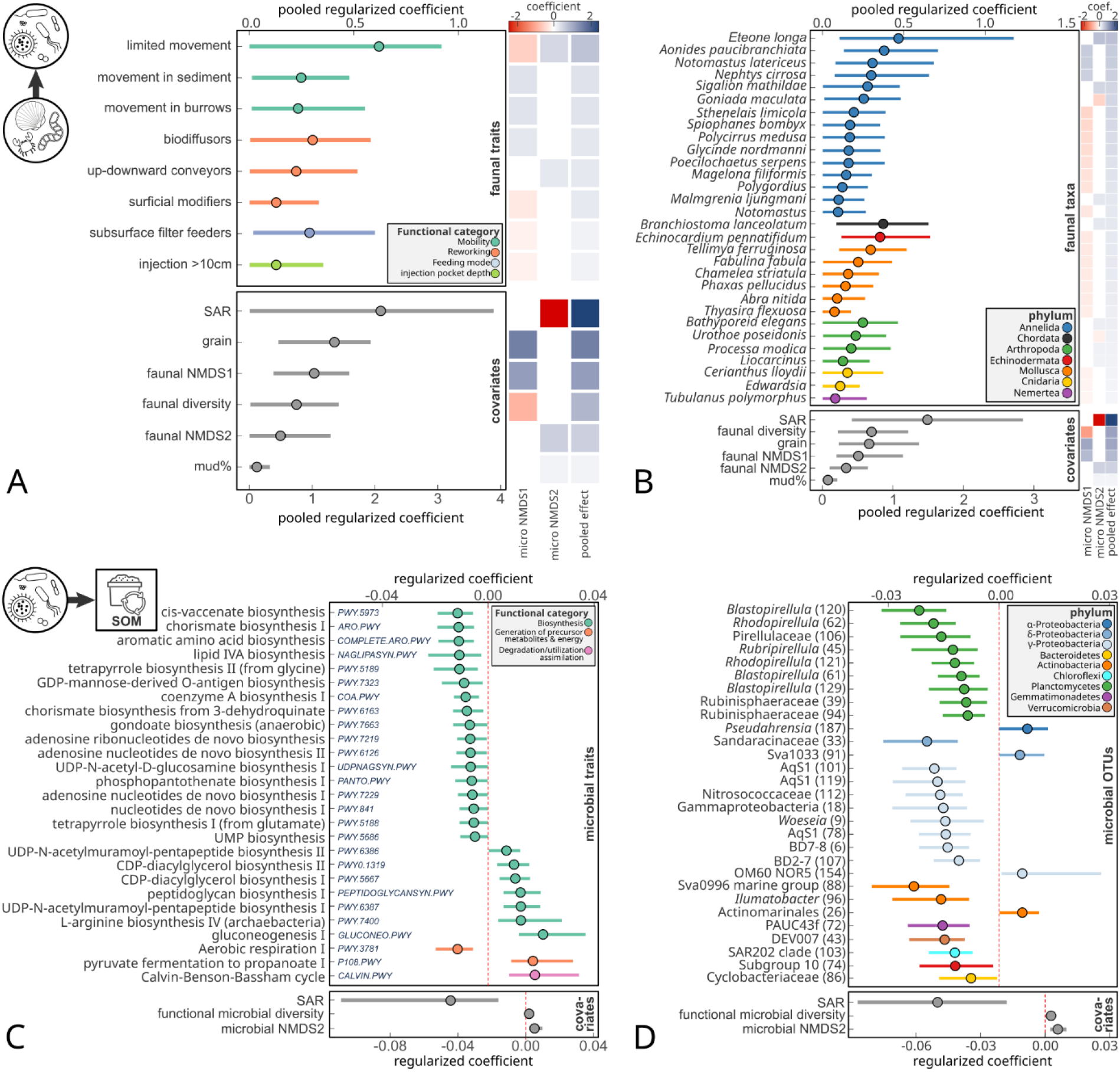
The most important faunal **(A, B)** and microbial **(C, D)** traits and species identified through pGLMs as key predictors of microbial community composition **(A, B)** and SOM% **(C, D). (A)** Pooled effect sizes of faunal traits and covariates retained from penalized regressions fitted on microbial NMDS1 and NMDS2. **(B)** Pooled effect sizes of faunal species and covariates retained from penalized regressions fitted on microbial NMDS1 and NMDS2. Pooled effects were calculated as the square root of the sum of the squared regularized effects of each predictor on NMDS1 and NMDS2. Regularized effect sizes from individual regressions are visualized in heatmaps adjacent to the plots. **(C)** Regularized effect sizes of microbial traits (e.g., pathways) and covariates retained from penalized regressions fitted on log-transformed SOM%. Pathway numbers are labeled within the plot area. **(D)** Regularized effect sizes of microbial OTUs and covariates retained from penalized regressions fitted on log-transformed SOM%. Colors correspond to phylum-level taxonomy.

Both the SEM and pGLMs consistently showed that changes in the microbial community were explained by macrofaunal diversity, suggesting that benthic microbiota are not only shaped by specific animal traits, but may also depend on ‘diversity effects’, such as resource partitioning, cooperation and facilitation (Fridley 2001; Gessner *et al*. 2010). Moreover, the positive indirect effect of macrofaunal diversity on SOM implies that diverse faunal communities mitigate microbial remineralization of SOM. In other words, faunal diversity favors OC storage. In marine sediments, studies on diversity effects are scarce, particularly in the context of interacting faunal and microbial communities. In terrestrial soils, complementarity processes (i.e., niche partitioning and functional differentiation among taxa) can increase OC assimilation into the food web and enhance OC decomposition (Gessner *et al*. 2010). Our results indicate similar processes in marine sediments. While this study focused on macrofauna and microbiota, meiofauna, an important component of benthic diversity and biomass (Gerlach 1971), were not considered. Meiofauna constitute a critical link between macrofauna and microbiota in the regulation of marine biogeochemical processes (Bonaglia *et al*. 2014). Accordingly, involving meiofauna in future studies will improve the mechanistic understanding of how the different components of the benthic biota interact and control carbon sequestration in marine sediments.

## CONCLUSIONS

The impacts of bottom trawling on SOM are not solely the result of direct physical disturbance, but also emerge from cascading effects involving benthic fauna and microbiota. Our results underscore the importance of ecological processes in trawling impacts on carbon sequestration, and demonstrate that benthic microbiota play a central role in seafloor carbon dynamics. Approximately one third of the trawling-associated decline in SOM is mediated by shifts in faunal and microbial communities, highlighting the need to consider biotic pathways in assessments of the impacts of bottom trawling on carbon sequestration. Considering these ecological processes in predictive models will be critical for more accurately forecasting the development of seafloor carbon stocks in the face of increasing anthropogenic disturbance. Future research should further disentangle how specific microbial and faunal functions interact with labile and recalcitrant OM to ultimately improve our understanding of seafloor carbon sequestration.

## MATERIALS AND METHODS

### Sampling and sediment properties

Macrofauna and microbiota were sampled with van Veen grabs (0.1 m^2^, minimum penetration depth: 10 cm) from 150 sites in the southeastern North Sea (Figure 1) with the German research vessel ‘Heincke’ in August 2019 (cruise HE538, doi:10.2312/cr_he538) and September 2020 (cruise HE562, doi:10.48433/cr_he562). At each site, one or three grabs were taken to characterize macrofaunal communities, and sediment samples were obtained from an additional grab taken to characterize microbial communities, grain size metrics, and SOM. The sediment temperature was measured on board using a standard thermometer (∼3 cm depth) from the first grab immediately after recovery. GPS coordinates were recorded from the ship-logs as the ship position at the time of bottom contact.

Samples for DNA extraction were taken from the top centimeter of the sediment directly in the grab (HE538) or with small vertical mini-cores (HE562), from which the upper centimeter was later sampled in the lab (details in Bonthond *et al*. 2023). A separate core (ø = 4.5 cm, penetration depth: 6 cm) was taken to characterize the sediment grain size distribution (Wentworth scale, Wentworth 1922) and total SOM content. Samples were stored at -20 °C until further processing. Subsamples of 40 g wet weight were dried and combusted for 5 hours at 500 °C to calculate SOM via loss on ignition (LOI) and characterize grain size distributions using a sieve cascade (details in Bonthond *et al*. 2023).

### Characterization of benthic microbiota

Benthic microbiota were characterized previously in Bonthond *et al*. (2023), where DNA extractions, library preparation, bioinformatic processing, OTU clustering (based on the 97% similarity criterion), and taxonomic classification are described in detail. For the present study, the final OTU table was rarefied by subsampling the count table a 100 times to a 1000 reads. Metagenomes were predicted with PICRUST2 v2.5.3 (Douglas *et al*. 2020), and MetaCyc pathway annotations were used for analyses.

### Characterization of benthic macrofauna and community properties

The van Veen grab contents were sieved (1 mm mesh size) onboard. Retrieved individuals of the benthic macrofauna were preserved in buffered 4 % formaldehyde-seawater solution and identified to the lowest taxonomic level possible in the laboratory and counted. The taxonomy was matched against the World Register of Marine Species (WoRMS Editorial Board 2024, https://www.marinespecies.org). The biomass was measured as fresh weight at the species level.

### Data preparation

To combine microbial and macrofaunal community data, datasets were reduced to a single observation per site. For sites with multiple mini-cores per sediment grab, the microbial community from the core taken from the most central position inside the grab was used. For sites with multiple macrofauna grabs, the grab nearest to the microbial grab was selected. As potential informative abiotic predictors for microbial and macrofaunal community composition and SOM, we considered the variables median grain size (µm), mud content (% dry weight), sediment temperature (°C), bottom shear stress (N/m^2^), and fishing intensity in terms of swept area ratio per year (SAR y^-1^). Fishing intensity was obtained from the OSPAR data & information management system (OSPAR 2017) and bottom shear stress was derived from the barotropic FESOM-C model (Androsov *et al*. 2019). Equivalent numbers of species, OTUs, and predicted MetaCyc pathways were used as a measure of faunal diversity, microbial diversity, and predicted functional microbial diversity, respectively (Jost 2006). The community bioturbation potential (BPc) was calculated according to Queirós *et al*. (2013), based on the biomass and abundance of macrofauna species, and species level scores for the traits ‘Mobility’ and ‘Sediment reworking’. For the community bioirrigation potential (IPc) we used the equation of Wrede *et al*. (2018), which uses the biomass and abundance of macrofauna species, and the traits ‘Burrowing’, ‘Feeding type’ and ‘Injection pocket depth’ (see Table S5 for more details on traits).

As general biotic predictors, we considered faunal biomass, faunal diversity, the BPc, IPc, and microbial diversity. Correlations among abiotic and biotic variables were explored using Spearman’s rank coefficients (Figure S1).

### NMDS to characterize macrofaunal and microbial community composition

Non-metric Multi-Dimensional Scaling (NMDS) was conducted with the R package *vegan* (v2.6-6.1, Oksanen 2010), using Aitchison distances for microbial communities (accounting for compositionality) and Bray-Curtis dissimilarities for macrofaunal communities. Abiotic variables and community properties were ordinated in NMDS spaces with the *enfvit()* function, evaluating significance with permutation tests with 9,999 permutations restricted within expeditions (Figure 2A, B).

### Linear mixed effect models (SEM pieces)

LMMs were fitted with restricted maximum Likelihood (REML) using the R-package *lme4* (v1.1-35.5, Bates *et al*. 2015), including the factor ‘Expedition’ as random intercept to account for the non-independence associated with expeditions. Moran’s I was computed to infer whether models accounted reasonably for spatial autocorrelation, considering autocorrelation negligible when |Moran’s I|< 0.1. To meet the model assumptions, we log-transformed the faunal biomass, BPc, IPc, and SOM. Variance inflation factors (VIFs) were computed to assess multicollinearity, and predictors with VIF values > 5 were excluded from LMMs.

Macrofaunal community structure was modeled by regressing both faunal latent variables against the abiotic predictors. Faunal biomass, faunal diversity, BPc, and IPc were fitted in response to the faunal latent variables. For the BPc and IPc, faunal biomass was also included as predictor. As we hypothesized that the impacts of abiotic variables on benthic microbiota are partially mediated by changes in macrofaunal communities, LMMs fitted on the microbial latent variables included the abiotic predictors, faunal biomass, BPc, IPc, faunal diversity, and the faunal latent variables. Based on variance inflation factor (VIF) scores, the faunal NMDS2 was excluded from both LMMs. Microbial diversity was regressed against the microbial latent variables, faunal diversity, and faunal latent variables. The final LMM, with SOM as response, included the microbial latent variables, representing microbial community composition, and faunal NMDS1 to represent microbial community composition. In addition, we included faunal and microbial diversity as predictors as they are known to influence OM cycling in soils (De Graaff *et al*. 2015; Gessner *et al*. 2010; Nielsen *et al*. 2011; Tao *et al*. 2023).

Further, faunal biomass, the BPc and the IPc were included as community level traits with potential impact on SOM as they may influence SOM positively (e.g., enhancing downward transport of OM) and negatively (e.g., faunal decomposition, upward transport and resuspension of SOM, Epstein *et al*. 2022). Finally, the model included the abiotic variables bottom shear stress and fishing intensity, which may reduce SOM through resuspension, potentially resulting in subsequent degradation of OM in the water column or transport to other areas (Epstein *et al*. 2022). Hence, temperature and sediment properties were only included as indirect effects on SOM in our SEMs, as they influence macrofaunal and microbial processes by regulating (heterotrophic) metabolic rates (Clarke & Fraser 2004) and the oxygen penetration depth (Neumann *et al*. 2021) but do not influence SOM directly.

### Structural Equation Models (SEMs)

We used structural equation modeling to disentangle the direct and indirect effects of abiotic and biotic variables on benthic microbiota and SOM. Faunal and microbial NMDS axes were used as latent variables, representing community structure. First, a conceptual causal network was constructed (Figure S2A) based on findings from previous work (Bonthond *et al*. 2023) and from literature (Gessner *et al*. 2010; Sciberras *et al*. 2018; Deng *et al*. 2020; Tiano *et al*. 2024). Using the framework of the *piecewiseSEM* R package (v2.3.0.1, Lefcheck 2016), LMMs were compiled into a SEM. After inspecting correlations, we included the following environmental variables: temperature, median grain size, mud content, bottom shear stress, and fishing effort. As it was found in Bonthond *et al*. (2023) that the median grain size and mud content can have opposite effects on microbial diversity and certain functions, it was decided to retain both variables in the analysis, even though they both represent sediment properties. In the first model, we included the faunal biomass, BPc, and IPc together, although these were strongly correlated (*ρ* > 0.75, Figure S1). Shipley’s test of d-separation revealed the presence of direct effects of temperature and trawling intensity on faunal diversity, which were added to the model (Figure S2). In addition, correlation terms were added to the global SEM between the IPc and BPc, microbial NMDS1 and NMDS2, and faunal NMDS2 and microbial NMDS2. As the faunal biomass and IPc were not informative as predictors, a reduced SEM was fitted without both responses, and both models were refitted with Maximum Likelihood and compared based on the corrected Akaike information criterion (AICc). The overall SEM fit was evaluated with Shipley’s test of d-separation obtained through Fisher’s C statistic and the significance of direct effects was assessed by bootstrapping the overall SEM with 9,999 iterations.

### Multivariate GLMs to identify trawling impacts on traits and taxa

To identify key traits and taxa responding to trawling effort, multivariate GLMs were fitted using the R package mvabund (v4.2.1), assuming a negative binomial distribution. First a model was fitted with the 100 relatively most abundant predicted pathways in response to the variables also considered in SEMs (temperature, median grain size, mud content, bottom shear stress and bottom trawling effort). P-values, indicating significant changes in relative OTU abundances in response to the predictors were obtained by resampling the model with 499 iterations with the *anova*.*manyglm* function, using ‘Expedition’ as blocking variable. The same model was fitted on the faunal traits and on OTUs and faunal taxa with > 0.1% relative abundance.

### Penalized GLMs to identify key traits and taxa

The effects of faunal traits and species on microbial community composition and microbial traits and OTUs on SOM were evaluated with separate pGLMs with trait category and species abundances as predictors, using the R package glmmLasso (v1.6.3). First, pGLMs were fitted on both microbial latent variables, using faunal trait categories related to bioturbation (‘Mobility’, ‘Reworking’, Queirós *et al*. 2013) and bioirrigation (‘Burrow type’, ‘Feeding type’, ‘Injection pocket depth’ – Wrede *et al*. 2018) as predictors, along with the environmental and community property covariates that were used in the SEM analysis. In a subsequent step, similar models were fitted where faunal traits were replaced by faunal taxon abundances. Second, a pGLM was fitted on SOM, using the microbial traits as predictors in combination with environmental variables and faunal community properties (faunal latent variables, BPc, faunal diversity) as covariates and microbial diversity and predicted functional microbial diversity. This step was repeated using the 100 most abundant OTUs as predictors instead of microbial traits.

In all pGLMs, faunal and microbial traits and species abundances were standardized by subtracting the mean and dividing by the standard deviation. A default lasso penalty of 10 was used to shrink coefficients toward zero in all models. The significance of selected predictors was inferred from 95% confidence intervals obtained by bootstrapping the models with 999 repeated iterations. Finally, for microbial community structure, we pooled the coefficients from both models (fitted on microbial NMDS1 and NMDS2) to estimate a single overall effect size for each predictor.

## Data Availability

Fastq files with raw de-multiplexed V3-V4-16S gene amplicon reads were generated in Bonthond *et al*. (2023) and are archived in the sequence read archive (SRA, Bioproject accession number PRJNA988469). The macrofauna data was deposited in CRITTERBASE (https://critterbase.awi.de; Teschke *et al*. 2022). Rendered R-scripts and respective input data for all analyses is available on GitHub at https://github.com/gbonthond/macrofauna-microbiota.

## Acknowledgements

The authors are grateful to the crew of the Research Vessel Heincke for support during both expeditions (HE538 in 2019, HE562 in 2020). Dr. Sahar Khodami and Dr. Pedro Martinez-Arbizu are thanked for amplicon sequencing at the Senckenberg am Meer Metabarcoding and SNG laboratory (this is publication number 112).

## Author contributions

GB, JB and LG conceptualized the study. Field collections were conducted by GB, JB, and LG. GB conducted laboratory work. GB, VS and JB processed data. GB conducted the formal analysis and drafted the manuscript. All authors contributed to writing and revising the manuscript.

