## Supplementary information for "Bottom trawling impacts on faunal and microbial communities increase seafloor carbon loss"

Benthic macrofauna and microbiota mediate bottom trawling impacts on long-term carbon storage in marine sediments

### SUPPLEMENTAL INFORMATION

**Table S1.** Permutation test for ordination vectors.

| | microbial NMDS | | | | | faunal NMDS | | | | |
|---|---|---|---|---|---|---|---|---|---|---|
| | NMDS1 | NMDS2 | R2 | Pr(>r) | | NMDS1 | NMDS2 | R2 | Pr(>r) | |
| SAR | -0.0992 | -0.99507 | 0.3821 | **0.0001** | *** | -0.20641 | -0.97847 | 0.2675 | **0.0001** | *** |
| temperature | 0.59952 | 0.80036 | 0.4247 | **0.0018** | ** | 0.921 | 0.38956 | 0.3665 | **0.0028** | ** |
| grain | 0.89736 | 0.4413 | 0.7997 | **0.0001** | *** | 0.99591 | 0.09033 | 0.6698 | **0.0001** | *** |
| SOM | -0.23346 | 0.97237 | 0.1956 | **0.0001** | *** | -0.36218 | 0.93211 | 0.1584 | **0.0001** | *** |
| mud | -0.78582 | 0.61846 | 0.3771 | **0.0001** | *** | -0.82386 | 0.56679 | 0.336 | **0.0001** | *** |
| shear | 0.97261 | 0.23246 | 0.3031 | 0.7751 | | 0.95712 | -0.2897 | 0.2994 | 0.3481 | |
| faunal mass | 0.49188 | 0.87066 | 0.3335 | **0.0001** | *** | 0.66741 | 0.74469 | 0.4832 | **0.0001** | *** |
| BPc | 0.23966 | 0.97086 | 0.3066 | **0.0001** | *** | 0.30583 | 0.95209 | 0.7781 | **0.0001** | *** |
| Ipc | 0.34256 | 0.9395 | 0.3162 | **0.0003** | *** | 0.49936 | 0.86639 | 0.5099 | **0.0001** | *** |
| faunal diversity | -0.68859 | -0.72515 | 0.2002 | **0.0389** | * | -0.46266 | -0.88654 | 0.4652 | **0.0001** | *** |
| microbial diversity | 0.99706 | 0.07668 | 0.182 | 0.2073 | | 0.66815 | -0.74402 | 0.2056 | **0.0429** | * |

*Note: The median gain size was transformed with a $log_2$. Faunal mass (Fmass), BPc, IPc and SOM were transformed with a natural logarithm. Correlation terms that were added to the SEM to satisfy Shipley's test of d-separation are indicated with tildes (~~).*

**Table S2.** Goodness of fit statistics of SEMs fitted in this study.

| SEM | Fisher' C | p value | DF | AIC | AICc |
|---|---|---|---|---|---|
| reduced SEM | 51.893 | 0.141 | 42 | 4507.120 | 4517.128 |
| large SEM | 80.238 | 0.189 | 70 | 5248.385 | 5261.859 |

**Table S3.** Output of the full SEM

| Response | Predictor | theoretical | | | | | | | bootstrapped | | | | | |
|---|---|---|---|---|---|---|---|---|---|---|---|---|---|---|
| | | Estimate | SE | DF | F | Std Est | p | | Std Est | bias | SE | $UCI_{95}$ | $UCI_{95}$ | |
| Mnmds1 | grain | 1.58 | 0.19 | 138.9 | 8.16 | 0.39 | <0.001 | *** | 0.24 | -0.02 | 0.03 | 0.20 | 0.32 | * |
| | temp | 0.14 | 0.15 | 62.7 | 0.93 | 0.04 | 0.354 | | 0.04 | -0.01 | 0.03 | -0.02 | 0.11 | |
| | mud | -0.31 | 0.16 | 138.2 | -1.91 | -0.06 | 0.059 | | -0.05 | 0.00 | 0.03 | -0.10 | 0.00 | * |
| | shear | 2.19 | 1.68 | 6.0 | 1.30 | 0.07 | 0.241 | | 0.05 | -0.02 | 0.04 | -0.03 | 0.13 | |
| | SAR | -0.45 | 0.81 | 23.6 | -0.56 | -0.02 | 0.583 | | -0.01 | 0.01 | 0.03 | -0.07 | 0.03 | |
| | Fmass | -0.06 | 0.21 | 136.2 | -0.28 | -0.01 | 0.779 | | -0.01 | 0.00 | 0.03 | -0.06 | 0.05 | |
| | BPc | -0.82 | 0.32 | 130.4 | -2.57 | -0.15 | 0.011 | * | -0.07 | 0.00 | 0.03 | -0.12 | -0.02 | * |
| | IPc | -0.08 | 0.24 | 138.1 | -0.33 | -0.02 | 0.741 | | -0.01 | 0.00 | 0.03 | -0.06 | 0.04 | |
| | Fdiversity | -1.11 | 0.33 | 123.9 | -3.40 | -0.13 | 0.001 | *** | -0.10 | 0.01 | 0.03 | -0.16 | -0.05 | * |
| | Fnmds1 | 1.87 | 0.18 | 136.6 | 10.53 | 0.55 | <0.001 | *** | 0.34 | -0.04 | 0.04 | 0.28 | 0.45 | * |
| Mnmds2 | grain | 0.61 | 0.18 | 138.3 | 3.33 | 0.34 | 0.001 | ** | 0.22 | -0.01 | 0.07 | 0.10 | 0.36 | * |
| | temp | -0.12 | 0.15 | 138.9 | -0.80 | -0.08 | 0.427 | | -0.07 | 0.01 | 0.09 | -0.25 | 0.10 | |
| | mud | 0.36 | 0.15 | 138.0 | 2.36 | 0.17 | 0.020 | * | 0.14 | 0.00 | 0.06 | 0.02 | 0.25 | * |
| | shear | -3.22 | 1.86 | 128.3 | -1.73 | -0.23 | 0.086 | | -0.17 | 0.02 | 0.10 | -0.40 | 0.01 | |
| | SAR | -1.97 | 0.83 | 136.9 | -2.37 | -0.21 | 0.019 | * | -0.14 | -0.01 | 0.06 | -0.25 | -0.01 | * |
| | Fmass | -0.26 | 0.20 | 138.5 | -1.31 | -0.15 | 0.191 | | -0.09 | 0.00 | 0.06 | -0.22 | 0.04 | |
| | BPc | 1.13 | 0.30 | 138.7 | 3.70 | 0.47 | <0.001 | *** | 0.22 | -0.01 | 0.06 | 0.13 | 0.35 | * |
| | IPc | 0.12 | 0.23 | 138.0 | 0.53 | 0.06 | 0.594 | | 0.03 | 0.00 | 0.06 | -0.09 | 0.15 | |
| | Fdiversity | 0.74 | 0.32 | 138.8 | 2.35 | 0.20 | 0.020 | * | 0.15 | -0.01 | 0.06 | 0.04 | 0.28 | * |
| | Fnmds1 | -0.72 | 0.17 | 138.5 | -4.25 | -0.49 | <0.001 | *** | -0.30 | 0.01 | 0.07 | -0.46 | -0.18 | * |
| Mdiversity | Mnmds1 | 5.29 | 2.34 | 143.0 | 2.27 | 0.37 | 0.025 | * | 0.18 | 0.00 | 0.08 | 0.04 | 0.34 | * |
| | Mnmds2 | -0.31 | 3.06 | 143.8 | -0.10 | -0.01 | 0.920 | | -0.01 | 0.00 | 0.07 | -0.15 | 0.13 | |
| | Fnmds1 | -13.56 | 7.97 | 143.5 | -1.70 | -0.28 | 0.091 | | -0.14 | 0.00 | 0.08 | -0.31 | 0.02 | |
| | Fnmds2 | -18.91 | 7.86 | 143.9 | -2.41 | -0.26 | 0.017 | * | -0.16 | 0.00 | 0.07 | -0.28 | -0.03 | * |
| | Fdiversity | 7.13 | 12.45 | 144.0 | 0.57 | 0.06 | 0.568 | | 0.04 | -0.01 | 0.07 | -0.09 | 0.20 | |
| Fnmds1 | grain | 0.76 | 0.07 | 143.1 | 10.90 | 0.63 | <0.001 | *** | 0.54 | 0.00 | 0.05 | 0.46 | 0.65 | * |
| | temp | 0.06 | 0.08 | 142.3 | 0.86 | 0.06 | 0.391 | | 0.06 | 0.01 | 0.07 | -0.08 | 0.18 | |
| | mud | -0.15 | 0.08 | 143.0 | -2.00 | -0.10 | 0.047 | * | -0.09 | 0.00 | 0.04 | -0.17 | 0.00 | * |
| | shear | -1.07 | 0.94 | 99.1 | -1.14 | -0.11 | 0.257 | | -0.08 | 0.02 | 0.08 | -0.28 | 0.05 | |
| | SAR | 0.87 | 0.40 | 131.8 | 2.21 | 0.14 | 0.029 | * | 0.10 | -0.01 | 0.05 | 0.01 | 0.21 | * |
| Fnmds2 | grain | 0.00 | 0.08 | 144.0 | 0.06 | 0.01 | 0.953 | | 0.00 | 0.00 | 0.08 | -0.15 | 0.16 | |
| | temp | 0.06 | 0.08 | 144.0 | 0.83 | 0.09 | 0.405 | | 0.06 | 0.01 | 0.08 | -0.12 | 0.21 | |
| | mud | 0.19 | 0.08 | 144.0 | 2.29 | 0.20 | 0.023 | * | 0.16 | 0.00 | 0.07 | 0.02 | 0.30 | * |
| | shear | -0.24 | 0.65 | 144.0 | -0.38 | -0.04 | 0.708 | | -0.03 | 0.00 | 0.10 | -0.23 | 0.18 | |
| | SAR | -1.79 | 0.36 | 144.0 | -4.97 | -0.43 | <0.001 | *** | -0.35 | 0.01 | 0.07 | -0.52 | -0.24 | * |
| Fdiversity | Fnmds1 | -0.11 | 0.03 | 144.8 | -4.00 | -0.28 | <0.001 | *** | -0.28 | 0.00 | 0.07 | -0.41 | -0.14 | * |
| | Fnmds2 | -0.39 | 0.04 | 144.1 | -10.45 | -0.65 | <0.001 | *** | -0.56 | 0.00 | 0.04 | -0.66 | -0.48 | * |
| | temp | 0.10 | 0.04 | 143.9 | 2.72 | 0.25 | 0.007 | ** | 0.24 | -0.01 | 0.09 | 0.08 | 0.42 | * |
| | SAR | -0.64 | 0.16 | 144.2 | -3.97 | -0.26 | <0.001 | *** | -0.22 | 0.00 | 0.06 | -0.34 | -0.12 | * |
| Fmass | Fnmds1 | 0.37 | 0.06 | 112.8 | 6.03 | 0.44 | <0.001 | *** | 0.44 | 0.02 | 0.07 | 0.28 | 0.57 | * |
| | Fnmds2 | 0.50 | 0.08 | 146.5 | 6.70 | 0.39 | <0.001 | *** | 0.39 | 0.00 | 0.05 | 0.28 | 0.49 | * |
| BPc | Fnmds1 | 0.10 | 0.02 | 146.0 | 4.21 | 0.17 | <0.001 | *** | 0.13 | 0.00 | 0.03 | 0.06 | 0.19 | * |
| | Fnmds2 | 0.59 | 0.03 | 146.0 | 17.67 | 0.63 | <0.001 | *** | 0.55 | 0.00 | 0.03 | 0.50 | 0.60 | * |
| | Fmass | 0.29 | 0.03 | 146.0 | 9.31 | 0.40 | <0.001 | *** | 0.29 | 0.01 | 0.03 | 0.22 | 0.34 | * |
| IPc | Fnmds1 | 0.07 | 0.04 | 16.9 | 1.75 | 0.10 | 0.099 | | 0.08 | 0.00 | 0.05 | -0.02 | 0.16 | |
| | Fnmds2 | 0.31 | 0.06 | 145.0 | 5.40 | 0.26 | <0.001 | *** | 0.23 | 0.00 | 0.04 | 0.15 | 0.31 | * |
| | Fmass | 0.60 | 0.05 | 115.3 | 11.26 | 0.66 | <0.001 | *** | 0.49 | 0.00 | 0.04 | 0.40 | 0.57 | * |
| SOM | SAR | -0.67 | 0.27 | 133.1 | -2.45 | -0.28 | 0.016 | * | -0.18 | 0.02 | 0.08 | -0.35 | -0.05 | * |
| | shear | 0.75 | 0.61 | 91.7 | 1.23 | 0.21 | 0.221 | | 0.15 | -0.04 | 0.13 | -0.06 | 0.46 | |
| | Fmass | -0.04 | 0.06 | 138.6 | -0.55 | -0.08 | 0.583 | | -0.04 | -0.01 | 0.08 | -0.20 | 0.13 | |
| | BPc | 0.05 | 0.10 | 138.6 | 0.46 | 0.08 | 0.646 | | 0.03 | 0.00 | 0.07 | -0.11 | 0.18 | |
| | IPc | 0.08 | 0.07 | 138.0 | 1.06 | 0.16 | 0.293 | | 0.08 | 0.00 | 0.08 | -0.06 | 0.23 | |
| | Fdiversity | 0.05 | 0.10 | 139.0 | 0.48 | 0.05 | 0.636 | | 0.04 | 0.00 | 0.08 | -0.11 | 0.18 | |
| | Mdiversity | 0.00 | 0.00 | 139.0 | 1.57 | 0.14 | 0.120 | | -0.02 | -0.01 | 0.09 | -0.18 | 0.16 | |
| | Fnmds1 | -0.02 | 0.08 | 139.0 | -0.26 | -0.05 | 0.797 | | 0.13 | -0.01 | 0.08 | -0.03 | 0.30 | |
| | Mnmds1 | 0.00 | 0.02 | 138.6 | -0.13 | -0.02 | 0.894 | | -0.01 | 0.01 | 0.08 | -0.18 | 0.16 | |
| | Mnmds2 | 0.10 | 0.03 | 138.9 | 3.61 | 0.37 | <0.001 | *** | 0.27 | -0.02 | 0.07 | 0.14 | 0.43 | * |
| ~~BPc | ~~IPc | 0.33 | | 150.0 | 4.27 | 0.33 | <0.001 | *** | | | | | | |
| ~~Mnmds1 | ~~Mnmds2 | -0.18 | | 150.0 | -2.15 | -0.18 | 0.016 | * | | | | | | |
| ~~Fnmds2 | ~~Mnmds2 | 0.14 | | 150.0 | 1.73 | 0.14 | 0.043 | * | | | | | | |

*Abbreviations: lower limit 95% confidence interval (LCI95), upper limit 95% confidence interval (UCI95). Note: The median gain size was $log_2$ transformed and Fmass, BPc, IPc and SOM were transformed with a natural log. Correlation terms added to the SEM to satisfy Shipley's test of d-separation are indicated with tildes (~~).*

**Table S4.** Output of the reduced SEM

| Response | Predictor | theoretical | | | | | | | bootstrapped | | | | | |
|---|---|---|---|---|---|---|---|---|---|---|---|---|---|---|
| | | Estimate | SE | DF | F | Std Est | p | | Std Est | bias | SE | UCI$_{95}$ | UCI$_{95}$ | |
| Mnmds1 | grain | 1.58 | 0.19 | 141.0 | 8.21 | 0.39 | **<0.0001** | *** | 0.24 | -0.02 | 0.03 | **0.20** | **0.32** | * |
| | temp | 0.16 | 0.15 | 85.8 | 1.05 | 0.05 | 0.296 | | 0.04 | -0.01 | 0.04 | -0.02 | 0.12 | |
| | mud | -0.30 | 0.16 | 140.0 | -1.87 | -0.06 | 0.064 | | -0.05 | 0.00 | 0.03 | -0.10 | 0.00 | |
| | shear | 2.27 | 1.71 | 8.9 | 1.33 | 0.07 | 0.218 | | 0.05 | -0.02 | 0.04 | -0.01 | 0.15 | |
| | SAR | -0.45 | 0.81 | 30.1 | -0.56 | -0.02 | 0.580 | | -0.01 | 0.01 | 0.03 | -0.08 | 0.04 | |
| | BPc | -0.96 | 0.21 | 141.0 | -4.63 | -0.17 | **<0.0001** | *** | -0.13 | 0.01 | 0.03 | **-0.18** | **-0.08** | * |
| | Fdiversity | -1.18 | 0.31 | 138.1 | -3.82 | -0.14 | **0.000** | *** | -0.11 | 0.01 | 0.03 | **-0.17** | **-0.06** | * |
| | Fnmds1 | 1.85 | 0.17 | 135.5 | 10.69 | 0.55 | **<0.0001** | *** | 0.34 | -0.04 | 0.05 | **0.29** | **0.47** | * |
| Mnmds2 | grain | 0.59 | 0.18 | 140.3 | 3.24 | 0.33 | **0.002** | ** | 0.21 | 0.00 | 0.07 | **0.09** | **0.35** | * |
| | temp | -0.09 | 0.15 | 140.9 | -0.62 | -0.06 | 0.533 | | -0.06 | 0.01 | 0.09 | -0.25 | 0.11 | |
| | mud | 0.37 | 0.15 | 140.0 | 2.48 | 0.17 | **0.014** | * | 0.15 | 0.00 | 0.06 | **0.03** | **0.26** | * |
| | shear | -3.02 | 1.85 | 126.8 | -1.63 | -0.22 | 0.105 | | -0.16 | 0.02 | 0.10 | -0.38 | 0.02 | |
| | SAR | -2.06 | 0.82 | 137.7 | -2.50 | -0.22 | **0.014** | * | -0.15 | -0.01 | 0.06 | **-0.26** | **-0.01** | * |
| | BPc | 0.98 | 0.20 | 140.3 | 4.94 | 0.41 | **<0.0001** | *** | 0.30 | -0.01 | 0.06 | **0.19** | **0.41** | * |
| | Fdiversity | 0.65 | 0.30 | 140.6 | 2.18 | 0.17 | **0.031** | * | 0.14 | -0.01 | 0.06 | **0.03** | **0.27** | * |
| | Fnmds1 | -0.75 | 0.17 | 140.7 | -4.47 | -0.51 | **<0.0001** | *** | -0.32 | 0.01 | 0.07 | **-0.46** | **-0.19** | * |
| Mdiversity | Mnmds1 | 5.29 | 2.34 | 143.0 | 2.27 | 0.37 | **0.025** | * | 0.18 | 0.00 | 0.08 | **0.03** | **0.34** | * |
| | Mnmds2 | -0.31 | 3.06 | 143.8 | -0.10 | -0.01 | 0.920 | | -0.01 | 0.00 | 0.07 | -0.15 | 0.13 | |
| | Fnmds1 | -13.56 | 7.97 | 143.5 | -1.70 | -0.28 | 0.091 | | -0.14 | 0.00 | 0.08 | -0.31 | 0.02 | |
| | Fnmds2 | -18.91 | 7.86 | 143.9 | -2.41 | -0.26 | **0.017** | * | -0.16 | 0.00 | 0.07 | **-0.28** | **-0.03** | * |
| | Fdiversity | 7.13 | 12.45 | 144.0 | 0.57 | 0.06 | 0.568 | | 0.04 | -0.01 | 0.07 | -0.09 | 0.20 | |
| Fnmds1 | grain | 0.76 | 0.07 | 143.1 | 10.90 | 0.63 | **<0.0001** | *** | 0.54 | 0.00 | 0.05 | **0.45** | **0.64** | * |
| | temp | 0.06 | 0.08 | 142.3 | 0.86 | 0.06 | 0.391 | | 0.06 | 0.01 | 0.07 | -0.08 | 0.18 | |
| | mud | -0.15 | 0.08 | 143.0 | -2.00 | -0.10 | **0.047** | * | -0.09 | 0.00 | 0.04 | **-0.18** | **0.00** | * |
| | shear | -1.07 | 0.94 | 99.1 | -1.14 | -0.11 | 0.257 | | -0.08 | 0.02 | 0.08 | -0.27 | 0.05 | |
| | SAR | 0.87 | 0.40 | 131.8 | 2.21 | 0.14 | **0.029** | * | 0.10 | -0.01 | 0.05 | **0.01** | **0.20** | * |
| Fnmds2 | grain | 0.00 | 0.08 | 144.0 | 0.06 | 0.01 | 0.953 | | 0.00 | 0.00 | 0.08 | -0.15 | 0.16 | |
| | temp | 0.06 | 0.08 | 144.0 | 0.83 | 0.09 | 0.405 | | 0.06 | 0.01 | 0.08 | -0.11 | 0.22 | |
| | mud | 0.19 | 0.08 | 144.0 | 2.29 | 0.20 | **0.023** | * | 0.16 | 0.00 | 0.07 | **0.02** | **0.30** | * |
| | shear | -0.24 | 0.65 | 144.0 | -0.38 | -0.04 | 0.708 | | -0.03 | 0.00 | 0.10 | -0.21 | 0.16 | |
| | SAR | -1.79 | 0.36 | 144.0 | -4.97 | -0.43 | **<0.0001** | *** | -0.35 | 0.01 | 0.07 | **-0.51** | **-0.22** | * |
| Fdiversity | Fnmds1 | -0.11 | 0.03 | 144.8 | -4.00 | -0.28 | **0.000** | *** | -0.28 | 0.00 | 0.07 | **-0.41** | **-0.14** | * |
| | Fnmds2 | -0.39 | 0.04 | 144.1 | -10.45 | -0.65 | **<0.0001** | *** | -0.56 | 0.00 | 0.04 | **-0.65** | **-0.48** | * |
| | temp | 0.10 | 0.04 | 143.9 | 2.72 | 0.25 | **0.007** | ** | 0.24 | -0.01 | 0.09 | **0.08** | **0.42** | * |
| | SAR | -0.64 | 0.16 | 144.2 | -3.97 | -0.26 | **0.000** | *** | -0.22 | 0.00 | 0.06 | **-0.34** | **-0.11** | * |
| BPc | Fnmds1 | 0.21 | 0.03 | 49.9 | 7.31 | 0.34 | **<0.0001** | *** | 0.34 | 0.02 | 0.05 | **0.23** | **0.42** | * |
| | Fnmds2 | 0.73 | 0.04 | 146.8 | 20.26 | 0.78 | **<0.0001** | *** | 0.78 | 0.00 | 0.02 | **0.74** | **0.83** | * |
| SOM | SAR | -0.70 | 0.27 | 135.5 | -2.57 | -0.29 | **0.011** | * | -0.19 | 0.02 | 0.08 | **-0.36** | **-0.05** | * |
| | shear | 0.80 | 0.60 | 96.2 | 1.32 | 0.22 | 0.190 | | 0.16 | -0.04 | 0.13 | -0.07 | 0.46 | |
| | BPc | 0.09 | 0.07 | 140.3 | 1.24 | 0.14 | 0.216 | | 0.09 | 0.00 | 0.07 | -0.05 | 0.24 | |
| | Fdiversity | 0.06 | 0.10 | 140.9 | 0.60 | 0.06 | 0.547 | | 0.05 | 0.00 | 0.08 | -0.11 | 0.19 | |
| | Mdiversity | 0.00 | 0.00 | 141.0 | 1.68 | 0.15 | 0.095 | | 0.14 | -0.01 | 0.08 | -0.02 | 0.31 | |
| | Fnmds1 | -0.01 | 0.07 | 141.0 | -0.19 | -0.04 | 0.847 | | -0.02 | -0.01 | 0.09 | -0.18 | 0.17 | |
| | Mnmds1 | 0.00 | 0.02 | 140.6 | -0.20 | -0.04 | 0.842 | | -0.02 | 0.01 | 0.08 | -0.19 | 0.14 | |
| | Mnmds2 | 0.10 | 0.03 | 141.0 | 3.68 | 0.37 | **0.000** | *** | 0.27 | -0.02 | 0.07 | **0.15** | **0.43** | * |
| ~~Mnmds1 | ~~Mnmds2 | -0.17 | | 150.0 | -2.12 | -0.17 | **0.018** | * | | | | | | |
| ~~Fnmds2 | ~~Mnmds2 | 0.15 | | 150.0 | 1.85 | 0.15 | **0.033** | * | | | | | | |

*Abbreviations: lower limit 95% confidence interval (LCI95), upper limit 95% confidence interval (UCI95). Note: The median gain size was $log_2$ transformed and Fmass, BPc, IPc and SOM were transformed with a natural log. Correlation terms added to the SEM to satisfy Shipley's test of d-separation are indicated with tildes (~~).*

**Table S5.** Traits and trait categories

| Effect trait | Trait Category | score | community trait index |
|---|---|---|---|
| Mobility[a] | organisms living in fixed tubes | 1 | BPc |
| | limited movement | 2 | |
| | free movement through sediment matrix | 3 | |
| | free movement via burrow system | 4 | |
| Reworking[a] | epifauna | 1 | |
| | surficial modifiers | 2 | |
| | upward and downward conveyors | 3 | |
| | Bio-diffusors | 4 | |
| | regenerators | 5 | |
| Burrow type[b] | epifauna or internal irrigation (e.g., siphons) | 1 | IPc |
| | open irrigation, (e.g., U or Y shaped burrows) | 2 | |
| | blind ended irrigation (e.g., blind end burrows, no burrow system) | 3 | |
| Feeding type[b] | surface filter feeder | 1 | |
| | predator | 2 | |
| | deposit feeder | 3 | |
| | subsurface filter feeder | 4 | |
| Injection pocket depth[b] | 0-2 cm | 1 | |
| | 2-5 cm | 2 | |
| | 5-10 cm | 3 | |
| | >10 cm | 4 | |

[a] Queirós et al. 2013. *Ecol. Evol.* 3: 3958–3985.
[b] Wrede et al. 2018. *Ecol. Indic.* 91: 737–743.

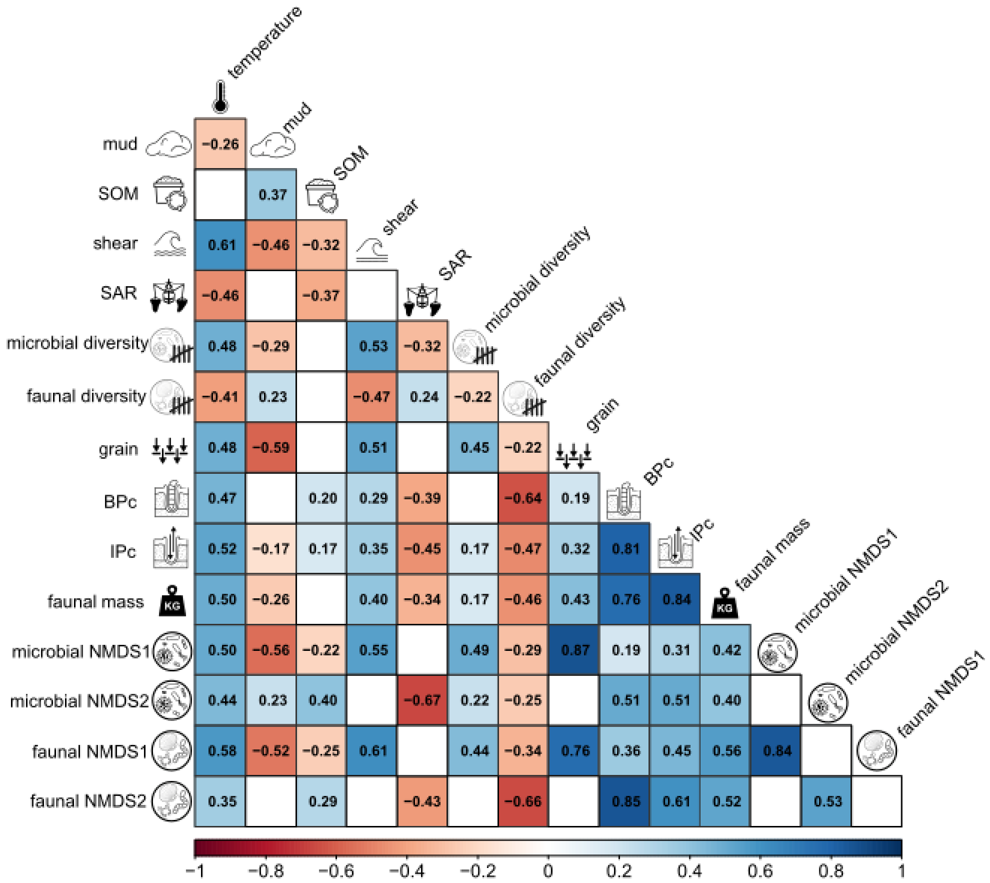


**Figure S1.** Matrix of Spearman's rank correlation coefficients among all predictor and response variables. The median grain size (grain) was transformed with a $\log_2$ and the community bioturbation potential (BPc), community bioirrigation potential (IPc), faunal mass and percentage sedimentary organic matter (SOM) with a natural log.

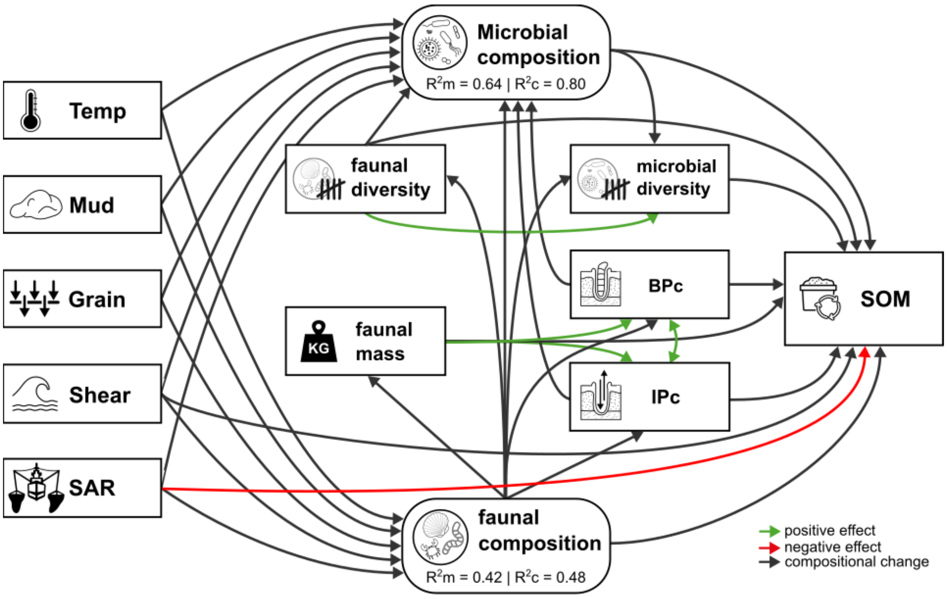


**Figure S2.** Hypothesized network of causal relationships. Squared boxes represent measured variables and rounded boxes pooled latent variables of faunal composition and microbial composition. For both faunal and microbial composition standardized effect sizes of both latent variables (NMDS axis 1 and NMDS axis 2) were pooled by taking the squared root of sum of the squared standardized effects).

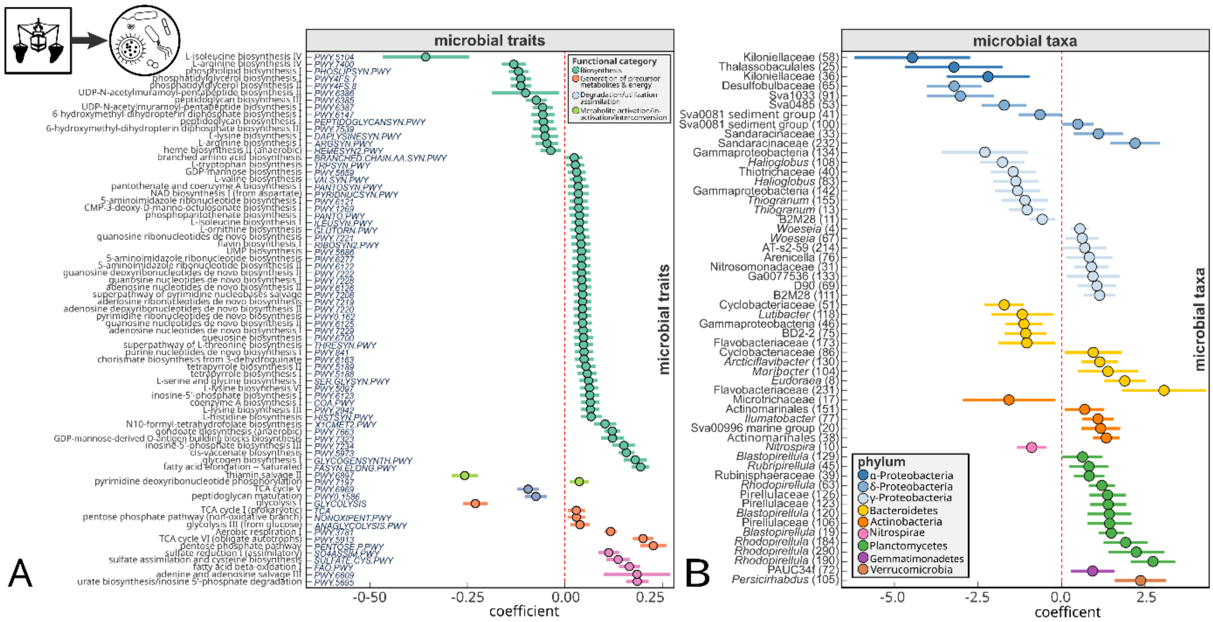


**Figure S3.** Trawling impacts on microbial traits and species identified through mGLMs (complete version of Figure 4A-B). **(A)** Microbial traits (predicted pathways) with significant responses to trawling effort. **(B)** Microbial taxa (OTU numbers in brackets) with significant responses to trawling effort.
